# Contrasting Patterns of Pierce’s Disease Risk in European Vineyards Under Global Warming

**DOI:** 10.1101/2023.07.17.549293

**Authors:** Àlex Giménez-Romero, Maialen Iturbide, Eduardo Moralejo, José M. Gutiérrez, Manuel A. Matías

## Abstract

Pierce’s Disease (PD) is a vector-borne disease caused by the bacterium *Xylella fastidiosa*, which poses a significant threat to grapevines worldwide. Despite its importance, the risk of future PD establishment in Europe remains unclear due to previous incomplete methodologies followed by conflicting results. Here we present a comprehensive approach considering the compound effect of climate change on the pathosystem. Within the general trend of progressively increasing PD risk, we identified the +3ºC scenario as a turning point for potential spreading beyond Mediterranean regions, representing a serious risk for French and Italian viticulture. Our innovative methodology reveals PD risk as a multi-factor multi-scale process, showing contrasting spatial patterns and different risk velocities across regions, as well as a high timing uncertainty. By overcoming previous limitations, our findings contribute to a better understanding of the potential spread of PD in Europe, supporting informed decision-making for disease management and prevention.

---

Climate change is widely recognized as an important driver of changes in the distribution and prevalence of plant diseases worldwide [1–6]. The impact of climate change on plant diseases has been approached from different perspectives [7, 8], but few studies have considered epidemiological dynamics in climate projections [9]. Modeling disease dynamics is a complex task, as they are emergent phenomena that arise from non-linear interactions among the disease components. In addition, many of the processes involved also show non-linear responses to changes in environmental variables [10, 11]. This complexity is further exacerbated in the case of vector-borne plant diseases (VBPDs) [12]. While climate primarily determines the potential geographic range of each organism in the pathosystem, the development of epidemic outbreaks depends on favorable host-pathogen-vector-climate interactions that drive transmission chains. Modeling the risk of emerging VBPDs implies delimiting their epidemiological niche rather than the ecological niche of their parts, as is commonly done. Therefore, when modeling risk in VBPDs it is essential to examine the complex spatial pattern that arises from the combined influence of climate on the distribution of hosts, pathogens and vectors.

Recently, a novel framework has been developed to properly evaluate the establishment risk of VBPDs [13]. It focuses on modeling the disease epidemiological niche based on the climate-driven spatial distribution of the vector, temperature-dependent bacterial growth and survival within hosts, and subsequent epidemiological dynamics. This has been specifically employed to predict the potential distribution of Pierce’s Disease (PD) worldwide. PD is a lethal disease of grapevines in the American continent that is transmitted unspecifically by sap-feeding insect vectors belonging to sharpshooter leafhoppers (Hemiptera: Cicadellinae) and spittlebugs (Hemiptera: superfamily Cercopidae) [14]. In the US, PD provokes huge economic losses to the wine sector estimated at 100 M$ per year in California alone [15]. The causative agent of PD is *Xylella fastidiosa* (Xf), a bacterium capable of colonizing the xylem vessels of more than 600 hosts, including important crops [16]. As a taxonomic unit, Xf comprises three recognized subspecies, *fastidiosa, multiplex*, and *pauca*, and more than 90 sequence types (i.e. genetic lineages) with different host ranges. Specifically, the Xf clonal lineage that causes Pierce’s disease (hereafter Xf_PD_) also causes almond leaf scorch disease in California [17].

Until the beginning of the 21st century, Xf was a pathogen officially restricted to the American continent [18]. In 2013, the involvement of Xf subsp. *pauca* in the massive death of ancient olive trees in Apulia, Italy, and its rapid spread raised alarm in European agriculture [19]. Today, all three Xf subspecies have been detected in the Balearic Islands (Spain), including Xf_PD_, and several clonal lineages have been found in Corsica and the PACA region of France, Alicante (Spain), Tuscany (Italy) and Portugal [20–22]. Outside the US, Xf_PD_ is only established on the islands of Mallorca and Taiwan, as well as recently detected in Israel, Lebanon and Portugal [23, 24]. In all European outbreaks, the insect vector *Philaenus spumarius* is the main and almost unique transmitter of Xf [25].

Recent studies indicate that the current risk of Pierce’s Disease (PD) establishment in Europe is primarily confined to the Mediterranean Basin [13, 26, 27]. Although there have been few efforts in characterizing the potential geographical distribution of Xf diseases in Europe under climate change scenarios, these attempts are limited to bioclimatic species distribution models (SDMs) of the pathogen [28–30] or the vector [27] using a few climate models. These independent analyses of two of the main disease components have led to conflicting results. Higher temperatures are expected to promote bacterial growth in susceptible crops in continental southern Europe, while these same areas will experiment with drier environmental conditions which are detrimental to vector populations [27]. In addition, the rather small number of climate models used in these studies makes it difficult to assess the uncertainty inherent in the predictions. These antagonistic patterns between the optimal ecological niche of the pathogen and its vector need to be urgently addressed to define the epidemiological niche of the disease and correctly assess the uncertainty in the prediction by using all available climate models.

Here we assess the potential distribution of PD under different global warming levels building upon the previous climate-driven epidemiological model [13]. To ensemble the epidemiological niche map, we used 40 of the most recent climate models and greenhouse gas emission estimates. Our study accounts for prediction uncertainties and provides an updated and comprehensive evaluation of the PD establishment risk in European wine-growing regions, addressing previous limitations. Additionally, it offers valuable insights for anticipating and managing the impacts of PD under a changing climate, which provides a basis for proactive measures to mitigate the risks associated with PD and ensure the resilience of viticulture despite future climate challenges.

## Present and future climate suitability of *Xylella fastidiosa* (PD) and *Philaenus spumarius*

To gain a deeper understanding of how climate change impacts each component of the pathosystem, we conducted separate analyses on the climatic suitability of Xf_PD_ and *P. spumarius*. Our model captures the thermal dependence of Xf_PD_ growth and survival within the infected vine through the ℱ(*MGDD*) and 𝒢(*CDD*) factors, respectively (see Methods). The suitability of Xf_PD_ infections is then determined by ℱ(*MGDD*) · 𝒢(*CDD*), providing the probability of symptom development during the growing season and subsequent survival for overwinter infection (see Methods). On the other hand, the climatic suitability of *P. spumarius* is modeled using a Species Distribution Model based on a previous study [27], where the indices CMI (Climatic Moisture Index; [31]) and TMAXSPRING (Mean maximum temperatures during the spring season) were identified as key predictors (see Methods).

Both analyses were evaluated under current (2003-2022) and future climate conditions considering increasing global warming scenarios (+1.5ºC, +2ºC, +3ºC, and +4ºC), obtained using the latest generation of regional climate projections over Europe [32] (see Methods). As the level of global warming increases, our model projects a substantial expansion of favorable conditions for Xf_PD_ in southern Europe with notable increments across the Mediterranean region (Fig. 1 and Supplementary Fig. S1), suggesting a potential spread of Xf_PD_ in these areas. Conversely, a decreasing trend in suitability was observed for *P. spumarius* in Southern Europe, although favorable conditions for the vector increased slightly in continental or hilly regions (Fig. 1 and Supplementary Fig. S1). In terms of absolute values, the increase in Xf_PD_ suitability across the different scenarios is remarkable, while the decrease in vector suitability is generally marginal, except in specific regions such as Spain and southern France (Supplementary Fig. S1).

**Figure 1:**
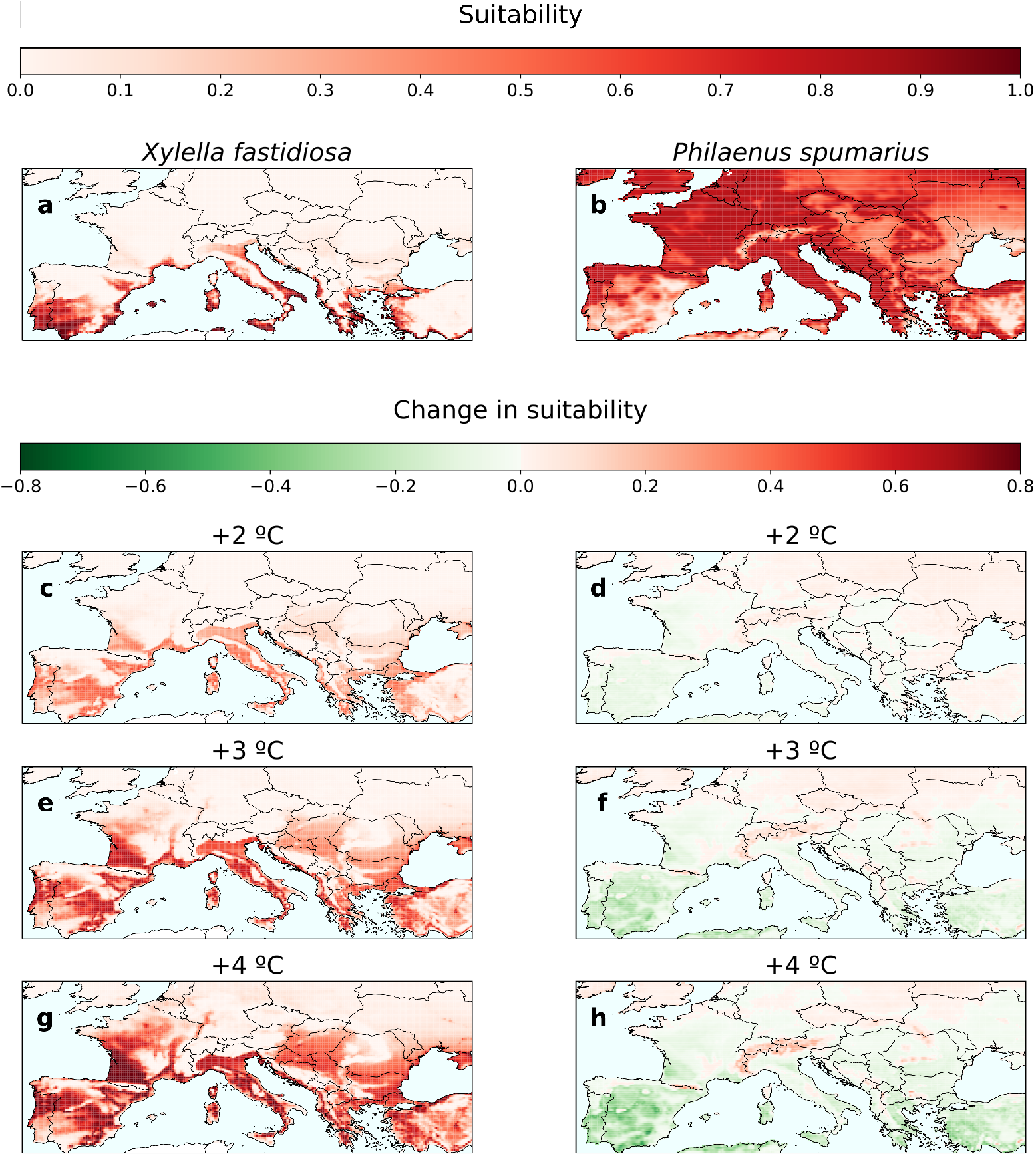
Change in Xf_PD_ and *P. spumarius* suitability under different climatic projections concerning the current scenario (2003-2022). (A, B) Current suitability. (C, D) +2ºC climatic projection. (E, F) +3ºC climatic projection. (G, H) +4ºC climatic projection. The suitability values for each scenario correspond to a 20 year average. The overall contrasting nature of expanding Xf_PD_ potential habitat and contracting *P. spumarius* ecological niche is clear.

An interesting observation is that a few new extensive suitable areas emerged suddenly, rather than expanding from previous ones. This phenomenon can be readily explained within our mechanistic framework. Western France lacks the protective winter-curing effect (𝒢 *>* 0.9), allowing climate change-induced increases in pathogen growth (from ℱ(*MGDD*) *<* 0.1 in a +1.5ºC scenario to ℱ(*MGDD*) *>* 0.6 in a +4ºC scenario) create a favorable habitat for Xf_PD_ (Supplementary Fig. S2). Conversely, Hungary and Serbia already experienced suitable conditions for pathogen growth in a +1.5ºC scenario (*ℱ*(*MGDD*) *>* 0.6), but were protected by relatively low survival rates (𝒢(*CDD*) *<* 0.3) (Supplementary Fig. S2). However, climate change would further boost pathogen growth and reduce the winter-curing effect, ultimately exposing the region to Xf_PD_ (Supplementary Fig. S2).

## The future risk of Pierce’s Disease in Europe

The contrasting dynamics of the Xf_PD_ and *P. spumarius* ecological niches give rise to fundamental questions regarding the establishment of PD in Europe. Can PD be established on the continent? Where and when it may occur? Our climate-driven epidemiological model directly focuses on delimiting the disease epidemiological niche to answer these questions. The model simulates an epidemic process in which the appearance of newly exposed hosts is influenced by the vector suitability while the transition to the infectious state is driven by the suitability of Xf_PD_ infections. The effective growth rate of the infected host population over the simulated period is used to derive a risk index *r*, bounded between − 1 and 1. Different risk categories naturally emerge under this modeling framework: no risk (*r* < − 0.1), transition zone (− 0.1 ≤ *r* < 0.1), low risk (0.1 ≤ *r* < 0.33), moderate risk (0.33 ≤ *r* < 0.66) and high risk (*r* ≥ 0.66). See the Methods section and the original work [13] for a detailed explanation of the model.

We found a general increase in the risk of PD across Mediterranean areas, with France, Italy and Portugal being particularly affected (Fig. 2 and Supplementary Fig. 2). Our findings uncover that a global temperature increase of +3ºC represents a turning point for the potential spread of PD beyond the Mediterranean region (Fig. 2 and Supplementary Fig. 2). Altogether, we found a general increase pattern for each of the designated risk categories across the different climate change scenarios (Supplementary Fig. S4).

**Figure 2:**
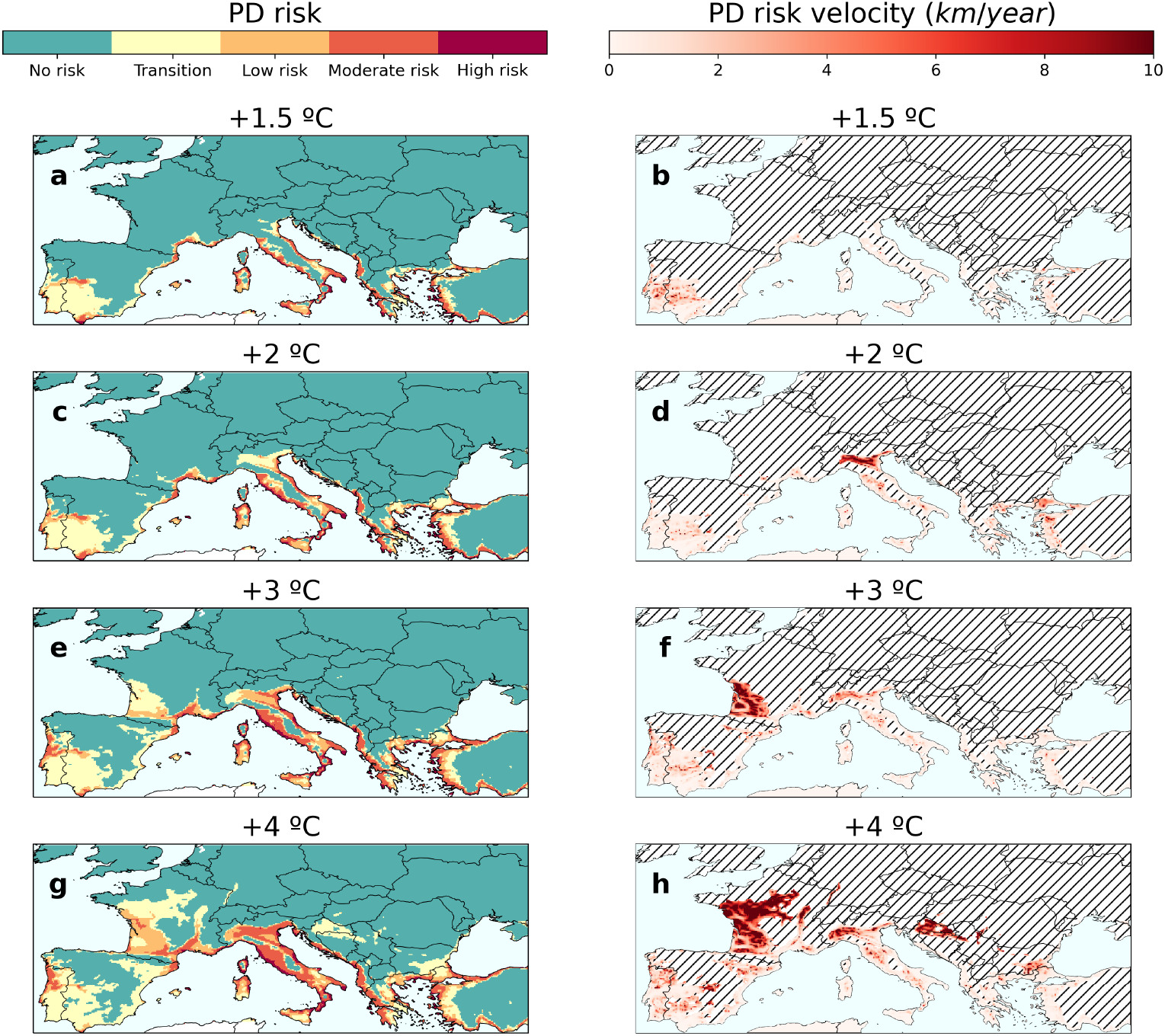
Risk of PD establishment and associated risk velocities under different climatic projections. (a,b) +1.5ºC climatic projection. (c,d) +2ºC climatic projection. (e,f) +3ºC climatic projection. (g,h) +4ºC climatic projection. Risk velocities were only computed in zones at risk in every scenario. Hatched lines in panels (b,d,f,h) indicate no risk zones where risk velocities have not been calculated.

To quantify the potential spread of PD, we computed the risk velocity, which allows to spot areas where the risk is changing rapidly or spreading at an accelerated pace (see Methods). We found a consistent and notable increase in the mean risk velocity within all designated risk zones, increasing from almost 1 km/year to 5 km/year as the temperature rises from a +1.5ºC to a +4ºC scenario (Supplementary Table S1). From the +2ºC scenario onward, PD risk velocities exhibit a substantial increase (Fig. 2 and Supplementary Fig. S4). At the +1.5ºC scenario, approximately 6% of the grid cells show risk velocities exceeding 5 km/year, whereas this value escalates to 50% under the +4ºC scenario (Supplementary Table S1). Moreover, a strong agreement between PD risk velocities and climate velocities was found, demonstrating that PD keeps trace with climate change [33].

These results represent the mean values obtained by averaging data from 40 climate models (see Methods). However, when considering the 16th and 84th percentiles (1*σ*), we observe significant variations in the predicted risk among different models. The results from the 84th percentile of a given warming level resemble those of the 16th percentile of a 2ºC higher warming scenario, indicating considerable uncertainty regarding the timing of occurrence Fig. 3. Nevertheless, the spatial patterns of risk are robust and consistent, indicating that while the timing may vary, the overall occurrence of the projected risks is likely to happen.

**Figure 3:**
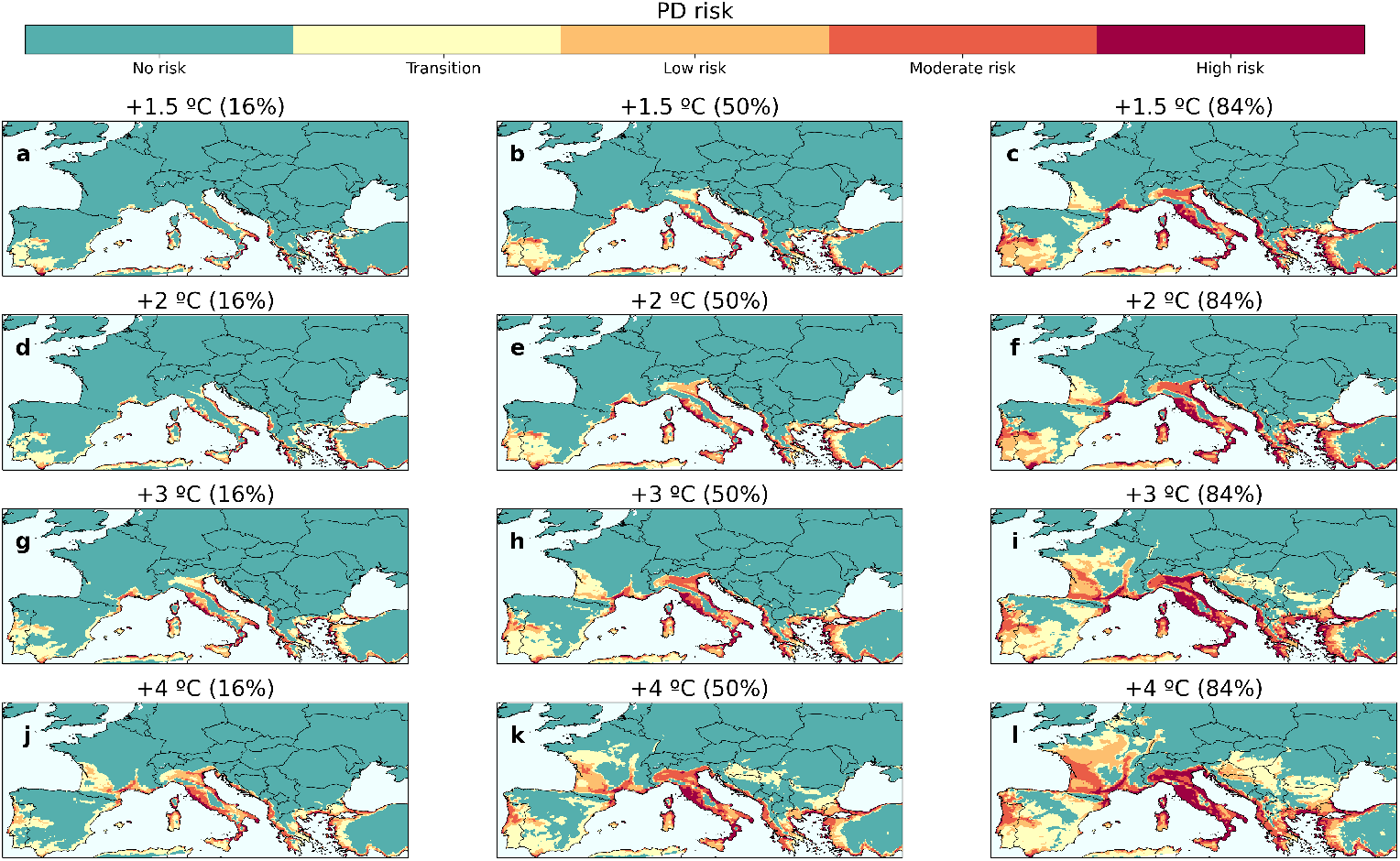
Uncertainty in PD risk projections under the different climate change scenarios. The estimates of PD risk under each climate scenario are given by the median risk values obtained from the 40 climate model ensemble (b, e, h, k). The uncertainty is obtained by considering the 1*σ* deviations from the central scenario, i.e. the 16% (a, d, g, j) and 84% (c, f, i, l) percentiles.

## The threat to European vineyards and Protected Designation of Origin regions

Accurately gauging PD risk and its effective impact requires consideration of the actual spatial distribution of vineyards. A comprehensive analysis is therefore needed across multiple scales, ranging from the country level to Protected Designation of Origin (PDO) regions and even down to individual vineyards.

Examining the proportion of the surface projected to be at risk on a country basis, we observe that Portugal and Greece face the highest overall risk. In a +1.5ºC scenario, these countries are projected to have 12% and 2% of their surface at risk, respectively, which escalates to a striking 47% and 63% in a +4ºC scenario. In contrast, countries such as France and Italy experience a lower, still relevant, increase in risk surface, never exceeding the 20% threshold, while Spain, the second largest wine producer, exhibits a decreasing trend of the risk areas above the +2ºC scenario Fig. 4 and Table 2. Such contrasting patterns in PD risk among countries only come to light when using our modeling framework.

**Figure 4:**
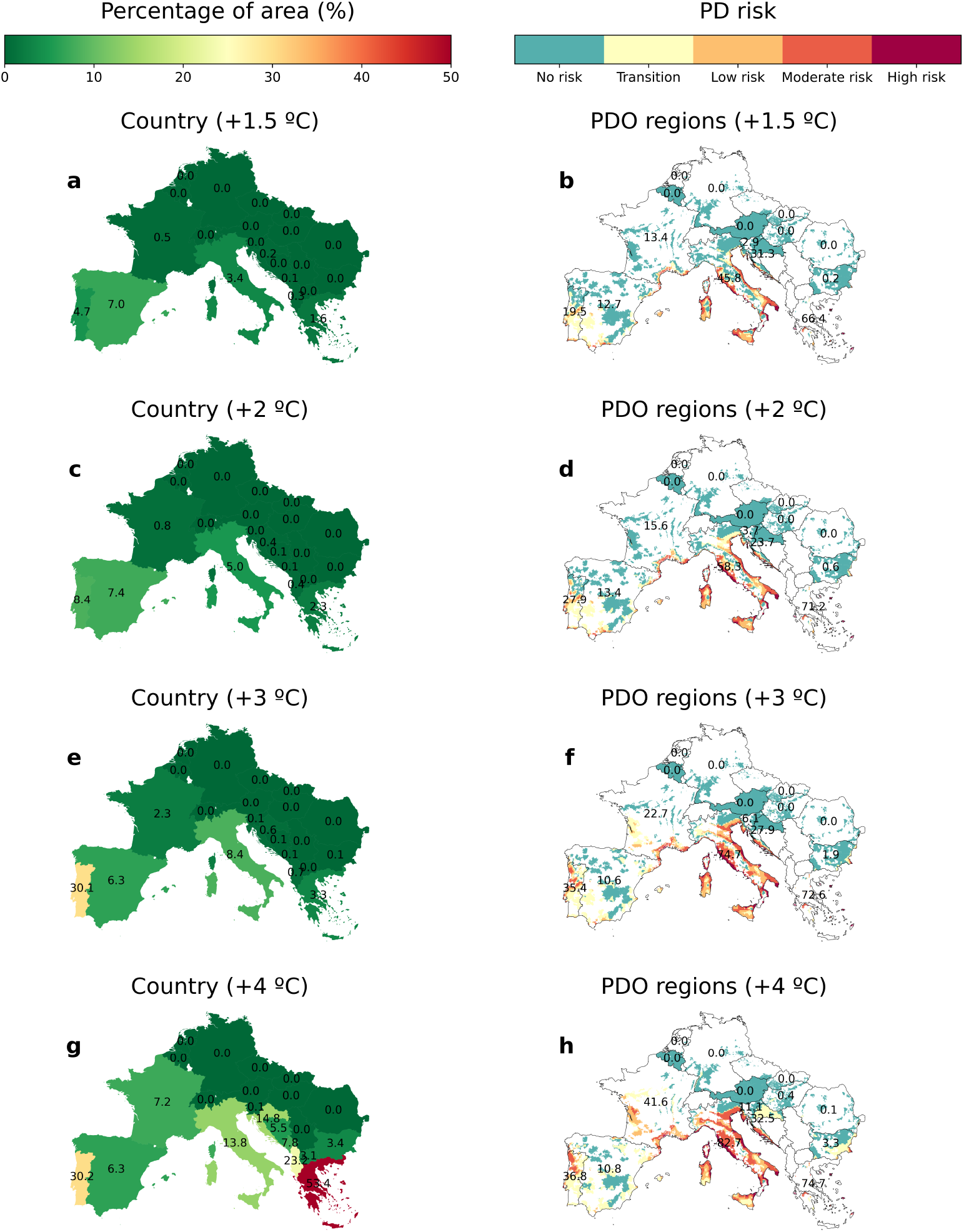
Multi-scale spatial analysis of PD future risk. (a,c,e,g) Percentage of country areas at risk (*r* > 0.1) for each climate projection. (b, d, f, h) Pierce’s Disease risk zones in Protected Designation of Origin wine regions for each climate projection. PDO data was obtained from [35]. The corresponding interactive analysis at the vineyard level can be found at [34]

A different picture is obtained when considering the spatial distribution of PDO regions and vineyards. For instance, PD risk within French and Italian PDO regions substantially increases from 13.4% and 45.8% in a +1.5ºC scenario to 41.6% and 82.7% in a +4ºC scenario, respectively Fig. 4 and Table 2, whereas the percentage of vineyard surface at risk surges from 24.21% and 57.49% in a +1.5ºC scenario to an astonishing 80% in a +4ºC scenario. Important European PDOs would be at risk from a +2ºC warming level, such as parts of the southern Rhône valley (Châteneuf du Pape), Provence and Languedoc in France, Penedés in Spain, Bairrada in Portugal, and Chianti and Brunello di Montalcino among others in Italy (see Supplementary Information). A detailed interactive analysis of the impact of PD in European PDO regions and vineyards is available on our web page [34].

Overall, our model simulations reveal a consistent increase trend in the risk of PD in Europe across all climate change scenarios. The percentage of the land surface at risk increases from 0.32% to 1.87%, the number of PDO regions at risk from 18.17% to 47.32%, and the vineyard surface from 18.67% to 40.35% Table 1.

**Table 1:** Percentage of land surface, vineyard surface and PDOs at risk in Europe under different climate projections.

| Scenario | Land surface (%) |  |  |  | Vineyard surface (%) |  |  |  | PDOs (%) |  |  |  |
| --- | --- | --- | --- | --- | --- | --- | --- | --- | --- | --- | --- | --- |
|  | +1.5 °C | + 2 °C | +3 °C | +4 °C | +1.5 °C | + 2 °C | +3 °C | +4 °C | +1.5 °C | + 2 °C | +3 °C | +4 °C |
| No risk | 99.4 | 99.15 | 98.61 | 96.14 | 68.89 | 61.3 | 45.54 | 37.64 | 72.7 | 64.88 | 49.17 | 33.89 |
| Transition | 0.28 | 0.39 | 0.64 | 1.99 | 12.44 | 14.35 | 21.86 | 22.0 | 9.13 | 9.39 | 15.28 | 18.79 |
| Low | 0.12 | 0.18 | 0.32 | 0.82 | 8.01 | 10.26 | 14.58 | 18.51 | 7.11 | 12.64 | 15.45 | 20.19 |
| Medium | 0.15 | 0.22 | 0.34 | 0.87 | 9.33 | 12.1 | 15.93 | 19.96 | 9.13 | 10.01 | 15.63 | 22.3 |
| High | 0.04 | 0.06 | 0.09 | 0.18 | 1.34 | 1.99 | 2.09 | 1.89 | 1.93 | 3.07 | 4.48 | 4.83 |
| Total risk (r>0.1) | 0.32 | 0.46 | 0.75 | 1.87 | 18.67 | 24.35 | 32.6 | 40.35 | 18.17 | 25.72 | 35.56 | 47.32 |

**Table 2:** Percentage of land surface, vineyard surface and PDOs at risk per different European countries under different climate projections.

| Scenario | Total surface (%) |  |  |  | Vineyard surface (%) |  |  |  | PDO (%) |  |  |  |
| --- | --- | --- | --- | --- | --- | --- | --- | --- | --- | --- | --- | --- |
|  | +1.5 °C | + 2 °C | +3 °C | +4 °C | +1.5 °C | + 2 °C | +3 °C | +4 °C | +1.5 °C | + 2 °C | +3 °C | +4 °C |
| Austria | 0.0 | 0.0 | 0.0 | 0.0 | 0.0 | 0.0 | 0.0 | 0.0 | 0.0 | 0.0 | 0.0 | 0.0 |
| Belgium | 0.0 | 0.0 | 0.0 | 0.0 | 0.0 | 0.0 | 0.0 | 0.0 | 0.0 | 0.0 | 0.0 | 0.0 |
| Bulgaria | 0.01 | 0.03 | 0.11 | 3.43 | 0.25 | 0.41 | 2.74 | 4.88 | 0.22 | 0.56 | 1.91 | 3.27 |
| Croatia | 0.24 | 0.38 | 0.64 | 14.82 | 31.53 | 42.6 | 45.89 | 47.65 | 31.27 | 23.75 | 27.93 | 32.47 |
| France | 0.53 | 0.84 | 2.32 | 7.25 | 24.21 | 35.75 | 56.43 | 80.0 | 13.37 | 15.62 | 22.73 | 41.65 |
| Germany | 0.0 | 0.0 | 0.0 | 0.0 | 0.0 | 0.0 | 0.0 | 0.0 | 0.0 | 0.0 | 0.0 | 0.0 |
| Greece | 1.62 | 2.33 | 3.32 | 53.37 | 59.61 | 65.08 | 70.36 | 66.69 | 66.38 | 71.19 | 72.61 | 74.68 |
| Italy | 3.44 | 4.98 | 8.39 | 13.8 | 57.49 | 65.4 | 77.34 | 81.79 | 45.77 | 58.34 | 74.71 | 82.65 |
| Portugal | 4.66 | 8.36 | 30.05 | 30.22 | 12.7 | 28.14 | 33.77 | 42.71 | 19.54 | 27.9 | 35.43 | 36.82 |
| Slovenia | 0.01 | 0.01 | 0.05 | 0.14 | 1.26 | 4.5 | 12.99 | 21.3 | 2.86 | 3.73 | 6.05 | 11.14 |
| Slovakia | 0.0 | 0.0 | 0.0 | 0.0 | 0.0 | 0.0 | 0.0 | 0.0 | 0.0 | 0.0 | 0.0 | 0.0 |
| Spain | 6.98 | 7.43 | 6.33 | 6.34 | 3.77 | 3.99 | 4.55 | 4.42 | 12.74 | 13.41 | 10.6 | 10.83 |

## Discussion

Previous research has made efforts to evaluate the potential geographic distribution of *Xylella fastidiosa* (Xf) and *Philaenus spumarius* in Europe using Species Distribution Models under future climates, reaching contrasting conclusions. While these suitability-based measures provide insights into the ecological niche of the disease’s key actors, their relationship with disease dynamics is still missing. Our approach integrates the compound effect of climate change in the pathosystem using a mechanistic epidemiological model to overcome these limitations. This innovative methodology has allowed us to address the important question about the potential future expansion of Pierce’s Disease (PD) in Europe. We must emphasize that although the model has been specifically applied to PD, the approach is rather general for vector-borne plant diseases.

One of the key findings of our study is the identification of a turning point for the risk of PD establishment at a +3ºC global mean temperature increase. Beyond this threshold, the risk of PD spreading north of the Mediterranean region becomes remarkably higher. This indicates that as global temperatures continue to rise, the range of PD could expand into new territories. Indeed, the projected increase in risk velocities under higher warming scenarios further emphasizes the potential for rapid spread and establishment of PD in previously unaffected regions. Regarding detailed potential impacts, Italian, French and Greek vineyards turn out to be the most vulnerable in Europe. However, it is crucial to acknowledge the uncertainty in this kind of prediction, which is not usually properly done. While earlier studies on the pathogen and vector distributions relied on a limited number of climate models, our results are based on an ensemble of 40 regional climate models, which reflect the most advanced knowledge available to date. This allowed us to observe a high variability in terms of the timing of occurrence, while still maintaining robust risk spatial patterns. In simple words, the spatial distribution of the establishment risk is clear, but it is uncertain exactly at which warming level this distribution would apply, always bounded by a ±1ºC increase.

Overall, our results highlight the contrasting effect of climate change on PD risk distribution in Europe, revealing it as a multi-factor and multi-scale process. Climate change has an opposite effect on each component of the pathosystem, enhancing areas of potential PD chronic infections while diminishing the suitable geographic range for the vector. At the same time, the characteristic spatial scale at which the risk is assessed highly influences conclusions. At the country level, there are significant variations in the extent of accumulated risk across different projections. However, when analyzed at a finer scale, such as at the level of PDO regions or vineyards, the results undergo a complete transformation. Countries that previously had marginal risk surfaces now show a higher percentage of PDO regions and vineyard surfaces at risk. These findings underscore the urgency of tailored mitigation and adaptation strategies to protect vineyards and PDOs, considering their specific spatial distribution and risk index, as well as the potential impacts of climate change.

It is important to acknowledge model limitations. Our results are influenced by the intrinsic uncertainty associated with correlative models used to determine the spatial distribution of the vector and the uncertainties in climatic projections. Although the spatial resolution in our climate projections is considered high, it may not capture the intricate micro-climate structure found in certain European wine-producing regions. Therefore, risk assessment results could locally differ with higher-resolution data. We did not include the possible influence of climate change on latitudinal and altitudinal shifts in the distribution of European viticulture [36, 37]. as this would only affect the calculation of the percentage of vineyard surface at risk but not the actual spatial distribution of risk. In any case, the risk estimates for the PDO regions include areas much larger than the areas of planted vines, which allows some margin in the adaptation and migration of the vineyards to different micro-climatic conditions. In addition, the PDO and vineyard databases used in this study are also subject to their own limitations. Future studies incorporating more refined modeling techniques and improved data resolution would enable a more nuanced understanding of PD risk and its potential impacts at the local scale.

Climate change is one of the biggest challenges for EU agricultural policy [38]. Quantitative regional predictions of climate change on emergent diseases like the present one provide a very valuable and unambiguous tool for decision-making. In our approach to the problem, risk indexes not only include information on where or where not PD can become established, but also reflect the exponential growth rate of potential epidemics, directly related to their potential economic impact. In addition, risk indices and velocities provide a dynamic framework for assessing the feasibility of eradication measures if Xf_PD_ is detected in a new area, offering crucial information for strategic crop protection. Our study evidences the need to selectively allocate more resources to surveillance and research on PD in countries of southern Europe, considering the associated uncertainties. This strategic allocation of resources based on risk assessment can help to prioritize proactive measures and effectively manage the potential impact of PD in different European countries.

In conclusion, our research highlights the complex dynamics of PD and its relationship with climate change. By adopting an interdisciplinary approach that integrates climate projections, epidemiological modeling, and spatial analysis, we provide valuable insights into the potential establishment and spread of PD in European wine-growing regions from country to vineyard levels. Our study demonstrates that accurately assessing the risk of PD establishment requires a nuanced understanding of the vector-plant-pathogen-climate system and the explicit consideration of the vineyard spatial setting. These findings can inform decision-making processes and support the development of effective strategies to mitigate the risks posed by PD and safeguard the future of viticulture in the face of a changing climate.

## Methods

### Climate datasets

We used E-OBS version v21e [39] as the reference observational climatic dataset for Europe, providing daily gridded data for Europe at a resolution of 0.1 degrees (∼10 km). In particular, maximum and minimum temperature data was used to compute the MGDD and CDD indices involved in the growth and survival processes of the Xf_PD_ pathogen (see “Climate-driven epidemiological model” section below).

To calibrate the distribution models of *Philaenus spumarius* capturing the widest possible range (North America and Europe), we used the ERA5-Land reanalysis [40]. This dataset was selected due to its global (land) coverage and high resolution (0.1 degrees, as E-OBS). In particular, daily precipitation and daily minimum and maximum temperature data were used to calculate the CMI and TMAXSPRING indices required for the vector suitability model (see “Vector suitability” section below).

Historical and future projections of the climate indices used in this study were calculated using regional climate simulations from the CORDEX initiative [41], in particular, the state-of-the-art large high-resolution (0.11 degrees) ensemble provided by EURO-CORDEX [32]. This dataset includes daily simulations of precipitation and temperatures from a large ensemble of Regional Climate Models (RCMs) driven by Global Climate Models (GCMs) from the CMIP5 project [42]. In particular, we considered the RCP8.5 simulations for 40 combinations of GCMs-RCMs (Table 3). In order to calculate 20-year mean climatologies of the indices across the different global warming levels (+1.5ºC, +2ºC, +3ºC and +4ºC), we relied on the time periods during which each CMIP5 driving model reaches the designated level within the RCP8.5 scenario (see [43]). This information is available at the IPCC WGI Atlas GitHub repository [44].

**Table 3:** EURO-CORDEX GCM-RCM combinations used in this study. Numbers indicate the number of runs in each combination.

|  | CNRM-CM5 | EC-EARTH | HadGEM2-ES | IPSL-CM5A-MR | MPI-ESM-LR | NorESM1-M |
| --- | --- | --- | --- | --- | --- | --- |
| CLMcom-CCLM4-8-17_v1 | 1 | 1 | 1 |  | 1 |  |
| DMI-HIRHAM5_v2 | 1 | 1 | 1 |  |  |  |
| GERICS-REMO2015_v2 | 1 |  |  |  |  |  |
| IPSL-WRF381P_v2 | 1 |  |  |  |  |  |
| KNMI-RACMO22E_v2 | 1 |  | 1 |  |  |  |
| SMHI-RCA4_v1 | 1 | 2 | 1 | 1 |  | 1 |
| CLMcom-ETH-COSMO-crCLIM-v1-1_v1 |  | 2 | 1 |  | 1 | 1 |
| DMI-HIRHAM5_v1 |  | 1 |  | 1 | 1 |  |
| IPSL-WRF381P_v1 |  | 1 | 1 | 1 |  | 1 |
| KNMI-RACMO22E_v1 |  | 2 |  | 1 | 1 | 1 |
| MOHC-HadREM3-GA7-05_v1 |  | 1 | 1 |  | 1 | 1 |
| GERICS-REMO2015_v1 |  |  |  | 1 |  | 1 |
| MPI-CSC-REMO2009_v1 |  |  |  |  | 1 |  |
| SMHI-RCA4_v1a |  |  |  |  | 1 |  |
| DMI-HIRHAM5_v3 |  |  |  |  |  | 1 |

### Climate-driven epidemiological model

We used the model developed in [13], which describes the initial exponential rise (or decrease) of infected plants at the onset of an epidemic based on two main features: the spatial distribution of the vector and the bacterial growth and survival processes mediated by temperature. In short, the density of vectors at a given site influences the number of new plants that will be inoculated with the bacterium, while the local temperature mediates the growth and survival processes of the in-plant bacterial population, leading to the initial inoculation to an infection or not. These temperature-driven growth and survival processes are described with the **accumulation** of two metrics denoted *Modified Growing Degree Days* (MGDD) and *Cold Degree Days* (CDD). The base function to compute the MGDD is proportional to the Xf temperature-dependent growth rate and is defined by

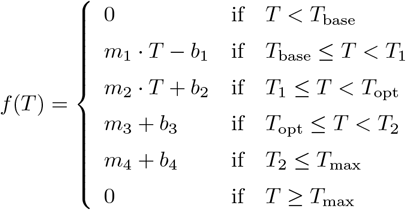

where *T*_base_ = 12 ^*◦*^C, *T*_1_ = 18, *T*_opt_ = 28 ^*◦*^C, *T*_2_ = 32 and *T*_max_ = 35 ^*◦*^C; the slopes are *m*_1_ = 0.66, *m*_2_ = 1, *m*_3_ = *−*1.25 and *m*_4_ = *−*3 and the intercepts are *b*_1_ =*−*8, *b*_2_ = *−*14, *b*_3_ = 4 and *b*_4_ = 105. MGDD are then computed between 1^st^ April and 31^st^ October as

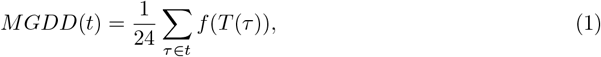

where *τ* is expressed in hours, *t* in years and we divide by 24 to obtain *MGDD*(*t*) in degree days. CDD are computed between 1^st^ November and 31^st^ March as

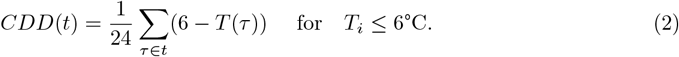

Altogether, the number of infected hosts is described by the following recurrence relation

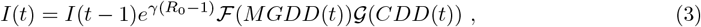

where *γ* is the death rate of infected vines, *R*_0_ is the basic reproduction number of the disease and ℱ(*·*) and *𝒢*(*·*) are sigmoidal-like functions that relate the MGDD and CDD metrics to the probability of developing an infection from a given inoculation. Following [13], *R*_0_ in each cell *j* is related to the climatic suitability of the vector such that

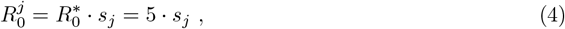

*γ* = 0.2 and the specific form of *ℱ*(*·*) and *𝒢* (*·*) is given by

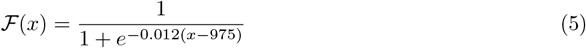

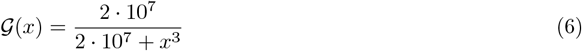

Finally, the risk index is derived as the effective growth rate of the infected population over the simulated time [13],

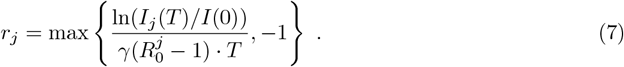

Because the typical time scale of the disease is 5 years (1*/γ*), we simulate periods of 7 years. If more years are available to simulate, we perform a re-introduction of the disease as a single infected plant in each cell after each 7-year period [13].

### Model adaptation to daily temperature data

The MGDD and CDD metrics used in the model were defined using hourly temperature data [13]. However, the E-OBS and CORDEX datasets only provide daily granularity. To overcome this limitation, we use a basic sinusoidal extrapolation relating maximum and minimum daily temperature to hourly temperatures,

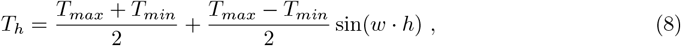

with *w* = 2*π/*24 and *h* ranging from 0 to 23.

This approximation was validated with data from national meteorological stations in Spain (AEMET) using several locations and years, showing that the computation of the MGDD and CDD metrics with hourly or daily temperatures yields the same results (Supplementary Fig. S5 and Supplementary Fig. S6).

Furthermore, we assessed the potential divergence of our temperature-based metrics (MGDD, CDD) when using ERA5-Land hourly mean temperature data (as done in [13]) compared to E-OBS daily maximum and minimum temperature data, used in this work. We must emphasize that not only the temporal resolution of the datasets are different, but also the data itself, which is acquired using different methodologies. Nevertheless, our results show that our metrics computed with both datasets are in good agreement with each other, showing a mean difference of 54 and 17 units for MGDD and CDD, respectively, and a standard deviation of 200 units for both metrics (Supplementary Fig. S7).

### Vector suitability

Following [45], we used the MaxEnt [46] algorithm to calibrate the relationship of *P. spumarius*global occurrence (predictand) with climate indices CMI and TMAXSPRING (predictors). Here we considered the most recent 20-year period with available data (2003-2022). The presence records of *P. spumarius* were obtained from The Global Biodiversity Information Facility (GBIF) [47][48]. Besides, we also used presence data collected by different Spanish plant protection agencies and research institutions (i.e. “Instituto de Ciencias Agrarias” at CSIC, Madrid, Spain; “Servicio de Sanidad Vegetal de la Junta de Andalucía” based in Sevilla and Jaén, Andalucía, Spain; Sanidad Agrícola Econex S.L. based in Murcia, Spain), as reported in [27]. A total of 1652 presence records were used (Fig. 5), ensuring that there were no duplicated records within each cell of the climate layer grid.

**Figure 5:**
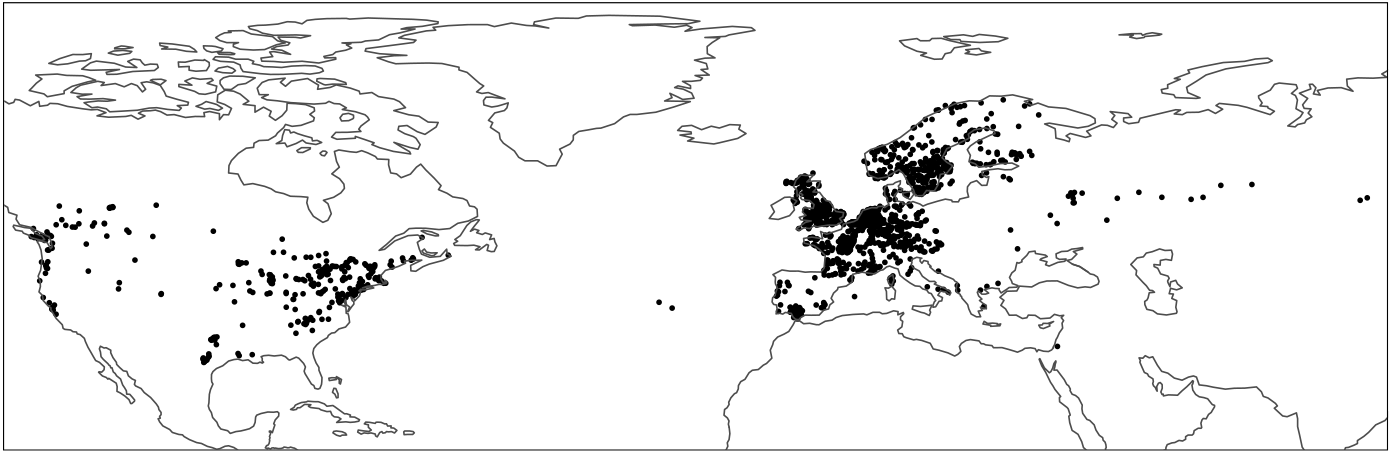
Training presence records for modeling the distribution of *Philaenus spumarius*.

In addition, we randomly generated pseudo-absences, also known as background points, using “The Three-Step” method proposed in [49]. This method incorporates a model performance criterion to determine the optimal sampling background extent, thereby ensuring that the model fitting was not adversely affected by the pseudo-absence sampling. Nevertheless, we accounted for the potential variability introduced by randomly selecting points from the background by performing 10 realizations of this sampling process. A total of 4956 pseudo-absences (three times the number of presences) were used in each realization.

Model evaluation was performed using a *k* -fold cross-validation approach (where *k* = 10) and the resulting AUCs (Area Under the ROC Curve) consistently exceeded 0.9 within the range of 0 to 1, with a value of 1 indicating perfect prediction and 0.5 indicating no discriminatory power (i.e. random guessing).

Finally, the calibrated models were used to predict the suitability of *P. spumarius* in the reference historical period (2003-2022) and under increasing global warming scenarios (panels b, d, f, h in Fig. 1).

### Risk velocity

To assess the dynamic nature of the risk index and its spatial propagation, we introduced the concept of risk velocity, a metric analogous to the recently proposed concept of climatic velocity [33]. The risk velocity represents the rate at which the risk index changes over time and spreads across different locations. Risk velocities were defined following the definition of climate velocity, as the ratio of the risk temporal trend and the risk spatial gradient in each cell. Thus, the units for the risk velocity correspond to kilometers per year (*km/*year). Risk velocities were computed using the VoCC R package [50, 51].

## Data availability

PDO regions georeference data was obtained from [35]. European vineyards georeferenced data was obtained from the Corine Land Cover [52]. MGDD, CDD and vector suitability data for the historical reference and the scenarios given by each warming level are available at [53]. We also provide the derived Xf_PD_ suitability, PD risk and PD risk velocity data in the same repository. The developed webpage is freely accessible at [34].

## Code availability

The code for the climate-driven epidemiological model is available in a GitHub repository [54]

## Acknowledgments

AGR and MAM were supported through grant PID2021-123723OB-C22 (CYCLE) funded by MCIN/AEI/10.13039/501100011033 and by “ERDF A way of making Europe” and through grant CEX2021-001164-M (María de Maeztu Program for Units of Excellence in R&D) funded by MCIN/AEI/10.13039/501100011033. MI and JMG acknowledge the E-OBS dataset from the EU-FP6 project UERRA (http://www.uerra.eu) and the data providers in the ECA&D project (https://www.ecad.eu). MI and JMG also acknowledge the World Climate Research Programme’s Working Group on Regional Climate, which is responsible for CORDEX, and we thank the climate modeling groups for producing and making available their model output. ERA5-Land [40] data was downloaded from the Copernicus Climate Change Service (C3S) Climate Data Store (2022) and neither the European Commission nor ECMWF is responsible for any use that may be made of the Copernicus information or data it contains. MI acknowledges support from project ATLAS (PID2019-111481RB-I00) funded by MCIN/AEI/10.13039/501100011033.

## Author contributions

AGR, MI, EM, JMG and MAM conceptualized the project and conducted investigations; MI processed the climate data; AGR performed the risk simulations and the analysis; AGR developed the webpage; AGR, MI, EM, JMG and MAM wrote the original draft; AGR, MI, EM, JMG and MAM reviewed and edited the manuscript; EM, JMG, MAM supervised the project; JMG and MAM acquired funding.

