## Supplementary Information for "Contrasting Patterns of Pierce’s Disease Risk in European Vineyards Under Global Warming"

### 1 Climate suitability for *Xylella fastidiosa* (PD) & *Philaenus spumarius*

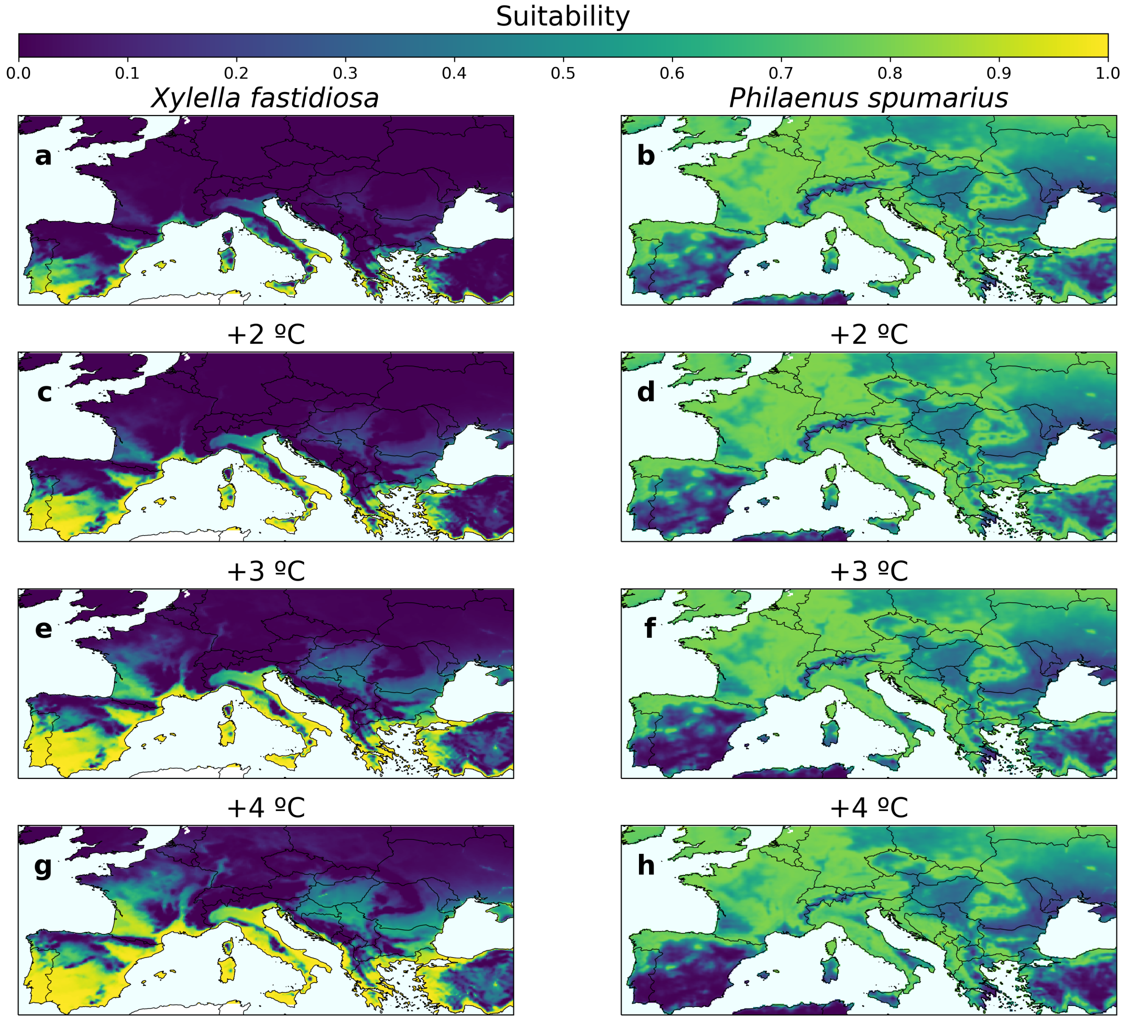


**Figure S1: Climate suitability of $Xf_{PD}$ and *P. spumarius* under the current scenario and different climate projections.** (a,b) Current scenario. (c,d) +2 ºC climate projection. (e,f) +3 ºC climate projection. (g,h) +4 ºC climate projection.

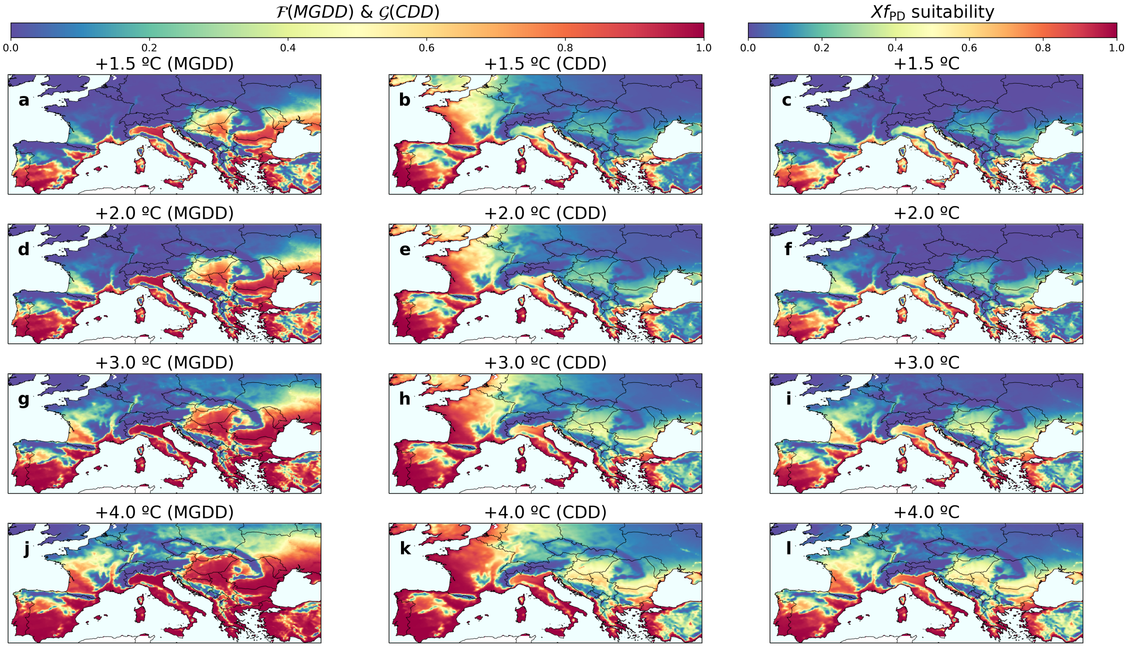


**Figure S2: $\mathcal{F}(MGDD)$, $\mathcal{G}(CDD)$ and suitability of $\mathbf{Xf_{PD}}$ ($\mathcal{F}(MGDD) \cdot \mathcal{G}(CDD)$) under different climate projections.** (a-c) +1.5 ºC climate projection. (d-f) +2 ºC climate projection. (g-i) +3 ºC climate projection. (j-l) +4 ºC climate projection.

### 2 PD future risk

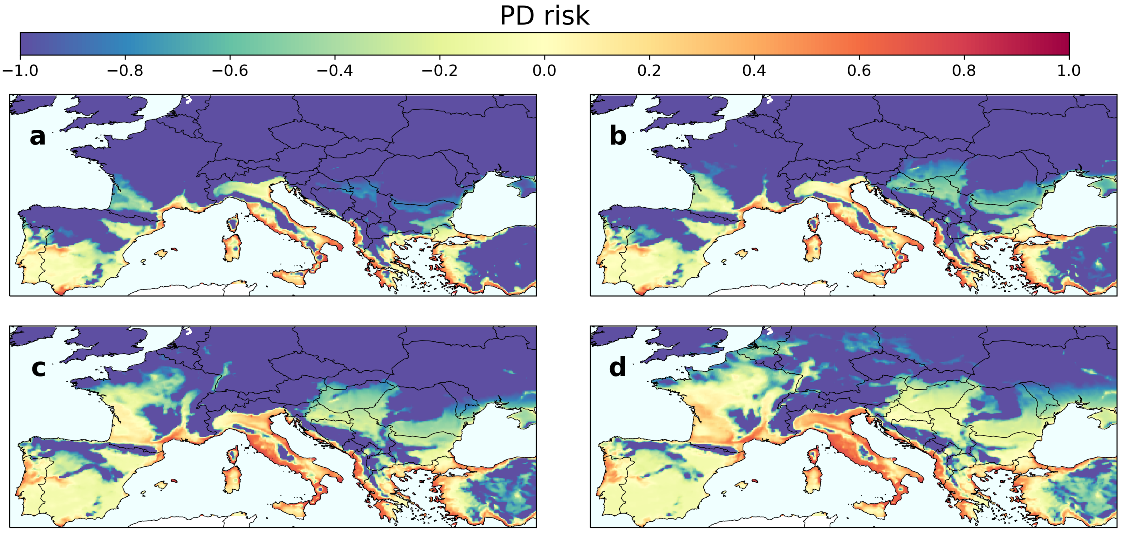


**Figure S3: Future risk of PD establishment under different climate projections.** (a) +1.5 ºC climate projection. (b) +2 ºC climate projection. (c) +3 ºC climate projection. (d) +4 ºC climate projection.

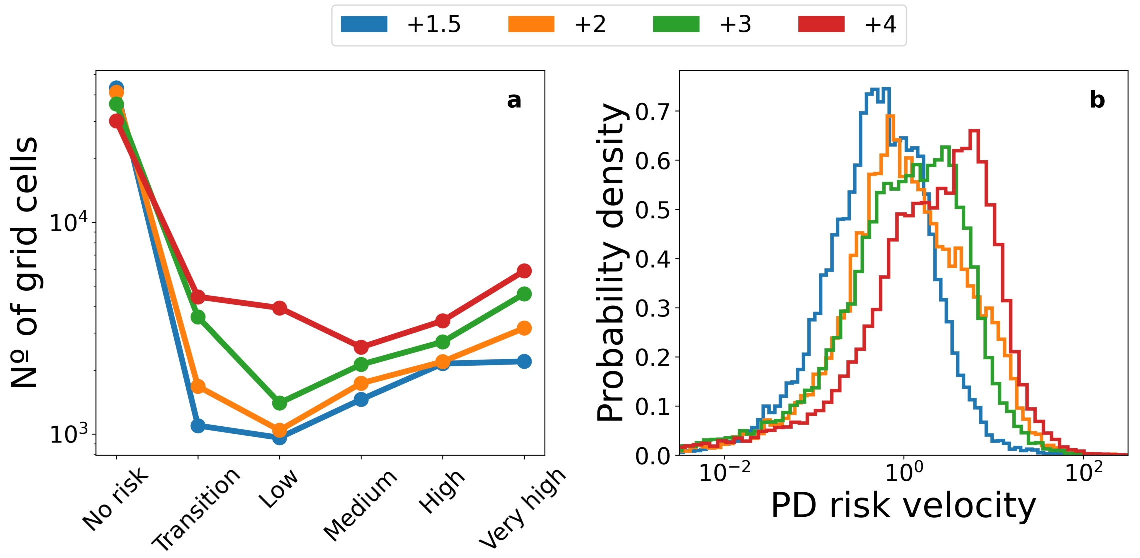


**Figure S4: Risk and risk velocity shifts as function of the climate projections.** (a) Number of grid cells at each risk level across the different climate projections. (b) Histograms of PD risk velocities in each climate projection. A shift toward higher velocities can be clearly observed.

**Table S1:** Some risk velocity statistics for each climate projection.

| **Scenario** | **+1.5 ºC** | **+2 ºC** | **+3 ºC** | **+4 ºC** |
|---|---|---|---|---|
| Risk velocity > 5 km/year (%) | 6.34 | 33.44 | 23.56 | 52.48 |
| Max risk velocity (km/year) | 181 | 691 | 393 | 667 |
| Mean risk velocity (km/year) | 0.78 | 1.9 | 3.84 | 5.15 |

### 3 Risk analysis within European wine Protected Designation of Origin regions

An extended analysis on the potential impact of PD in European PDOs can be easily performed using our PD establishment risk webpage, where the following insights can be contrasted in an interactive way.

Some French wine regions consistently remain risk-free in all scenarios, such as Alsace, Jura, and Champagne, while others, like Bourgogne (including Beaujolais) and the Loire Valley, only reach the transitional risk region in the warmest scenarios . Conversely, the Rhône Valley, Southwest, Languedoc, Roussillon, Provence, and Bordeaux face increasing risk in warmer scenarios. Within the Rhône region, Condrieu PDO stays out of risk in the northern part, while Côte-Rotie reaches low risk only in the +4 ºC scenario. Further south, Châteauneuf-du-Pape reaches low risk from the +2 ºC scenario, and Costières de Nîmes becomes low risk from the +1.5 ºC scenario, escalating to medium risk from the +2 ºC scenario. Regions directly influenced by the Mediterranean, like Cassis and Bandol in Provence or Muscat de Frontignan in Languedoc, experience higher risk, with some coastal areas reaching medium-high risk levels. In Provence, the risk becomes medium in the warmest scenarios, while in Languedoc-Roussillon, it typically decreases to low risk from the +2 or +3 ºC scenarios (except for Muscat DOPs). In Bordeaux, certain right bank PDOs (Pomerol, Lalande de Pomerol) become low risk in the +3 ºC scenario, while Saint Emilion achieves the same in the +4 ºC scenario. On the left bank, major DOPs such as Margaux, Saint Julien, Pauillac, and Saint Estèphe reach low risk in the +4 ºC scenario, along with Pessac-Léognan in the region below the city of Bordeaux. Graves and Sauternes, however, reach at most the transitional risk level in the +3 ºC scenario.

In Spain, contrasting patterns are observed. The northwestern part, including Rioja and Ribera del Duero DOPs, consistently remains risk-free in all four scenarios, except for the coastal Rías Baixas DOP, which becomes low risk in the +4 ºC scenario. In the southern region of Andalusia, a decrease in risk level is observed with increasing warming, as seen in the cases of the Jerez/Sherry PDO (labeled as Manzanilla) and Sierras de Málaga, which have low risk in the +1.5 ºC and +2 ºC scenarios and become transitional at +3 ºC and +4 ºC, respectively. This decrease in risk is associated with decreased vector suitability as temperatures rise. In the more continental DOPs of Aragón and Valencia (Utiel-Requena), the risk remains non-existent, while those closer to the Mediterranean exhibit increased risk levels. Penedès becomes low risk in the +2 ºC scenario, and the coastal DOPs of Alella and Empordà clearly increase to medium risk at the +2 ºC scenario from a low risk level at the +1.5 ºC scenario. Other more continental DOPs, such as Priorat, become transitional at most. A similar pattern is observed in central Spain, with northern DOPs remaining risk-free and more southern ones (Méntrida, Ribera del Guadiana) becoming transitional. Notably, the large La Mancha DOP remains risk-free, again due to decreased vector suitability.

In Portugal, a similar pattern of decreasing risk is observed in the southern regions of Algarve and Setúbal. However, there is a clear increase in risk in the central DOPs of Bairrada (reaching low-risk in the +2ºC scenario and medium-risk from +3ºC), Dão DOP (becoming low-risk from the +3ºC scenario), and Vinho Verde DOP, which exhibits the same pattern as the Rías Baixas region in Galicia, becoming low risk in the +4ºC scenario. The Óbidos DOP also becomes low risk from the +2ºC scenario. The Carcavelos and Colares DOPs, with their strong maritime influence, maintain or reach a medium risk level, while other DOPs like Alentejo, Beira Interior, and Trás-os-Montes are not at risk. The Douro DOP transitions from no risk to the transitional level, but this change may be influenced by the relatively low spatial resolution of our study, as the wine-producing area is relatively narrow (less than 0.1º).

In Italy, there is an overall increase in risk observed across the country. However, the increase is more limited in Piedmont, where DOPs such as Barolo, Alba, Barbera, and Langhe only reach low-risk levels at 4ºC. Similarly, in Veneto and Friuli, they become low risk drom the 3ºC scenario, while Alto Adige DOP remains non-risk. In Tuscany, there is a considerable higher increase in risk. Coastal regions like Maremma and Bolgheri reach medium and high risk at 2ºC, respectively, having positive risk already in the 1.5ºC scenario and the island of Elba already is already at high risk in the base scenario. More interior tuscan DOPs like Chianti and Brunello di Montalcino become low-risk from the +2ºC scenario. In Umbria, the risk increases rapidly, with Colli Perugini DOP reaching medium-risk at the +3ºC scenario, and Torgiano and Orvieto transitioning from low-risk at +2ºC to medium risk at +3ºC. In the Apulia region, there are distinct differences between the lower tip of the Apulian peninsula, where the risk levels remain consistently high, and the more continental north, where DOPs like Gravina in Bari province become low-risk from the +2ºC scenario, and San Severo DOP in Foggia province consistently stays at low-risk across all scenarios. In the islands, the risk decreases with increasing warming scenarios in some zones of Sicily, like Marsala, while others stay at risk and not changing the level, and a nonmonotonic behavior is found in Etna (decreases at 2ºC and increases again at 3ºC), while Sardinian DOPs maintain their risk level, typically medium, like Malvasia di Bosa, Vermentino di Gallura and Vernaccia di Oristano.

### 4 MGDD and CDD from daily temperature approximation

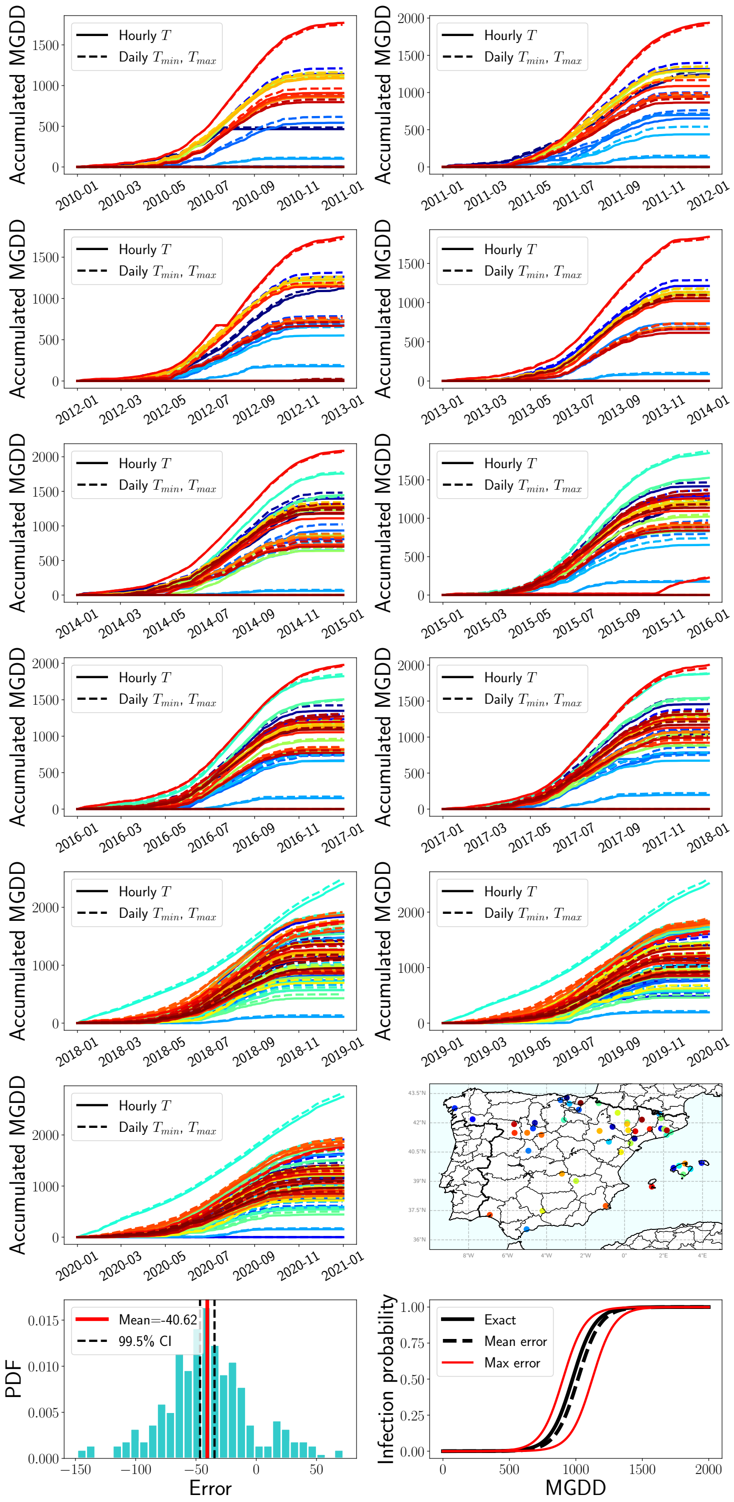


**Figure S5:** Comparison of MGDD accumulation computed with hourly mean temperature data (solid line) and daily maximum and minimum values (dashed line) using data from several meteorological stations in Spain and different years. The last row shows the distribution of errors and mean error in MGDD and the mean and maximum potential errors in infection probability.

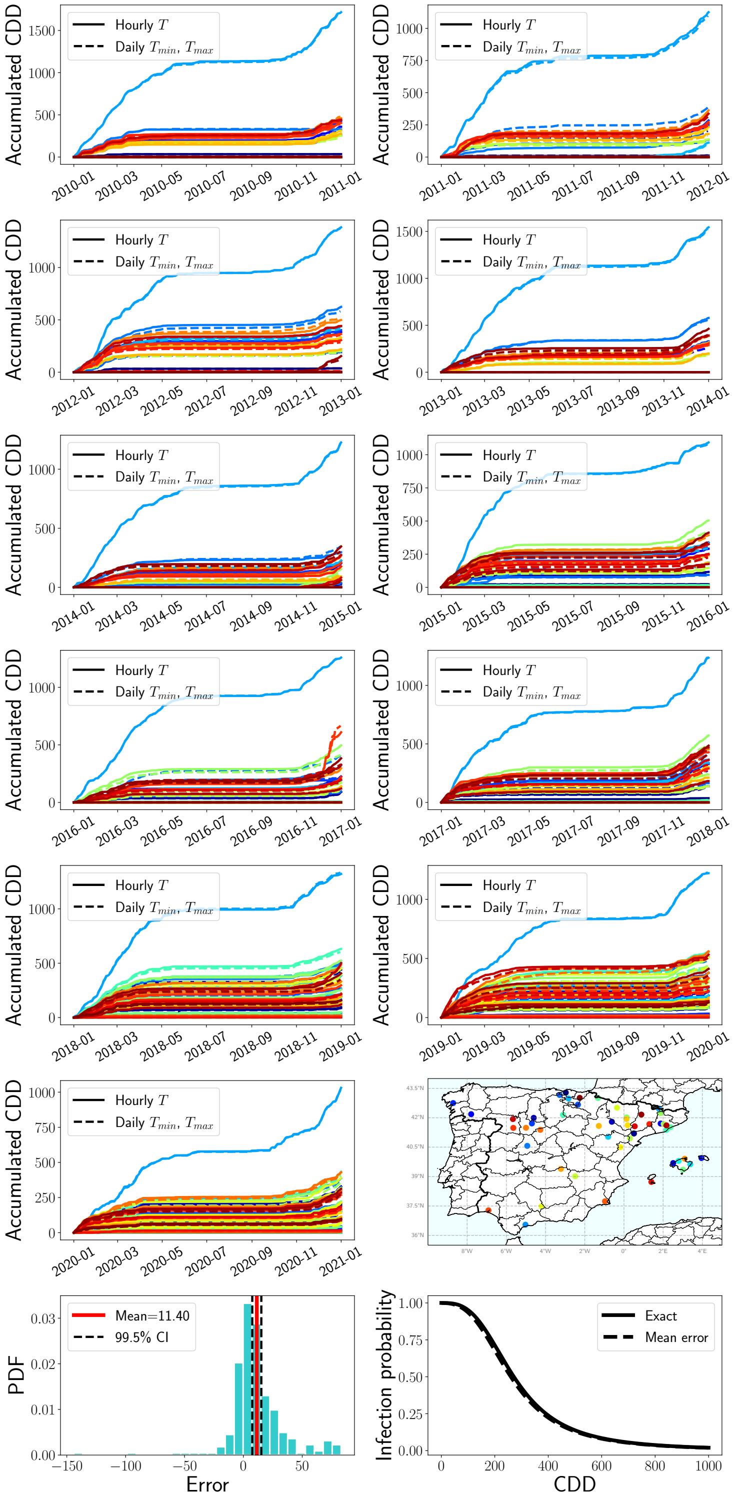


**Figure S6:** Comparison of CDD accumulation computed with hourly mean temperature data (solid line) and daily maximum and minimum values (dashed line) using data from several meteorological stations in Spain and different years. The last row shows the distribution of errors and mean error in CDD and the mean and maximum potential errors in infection probability.

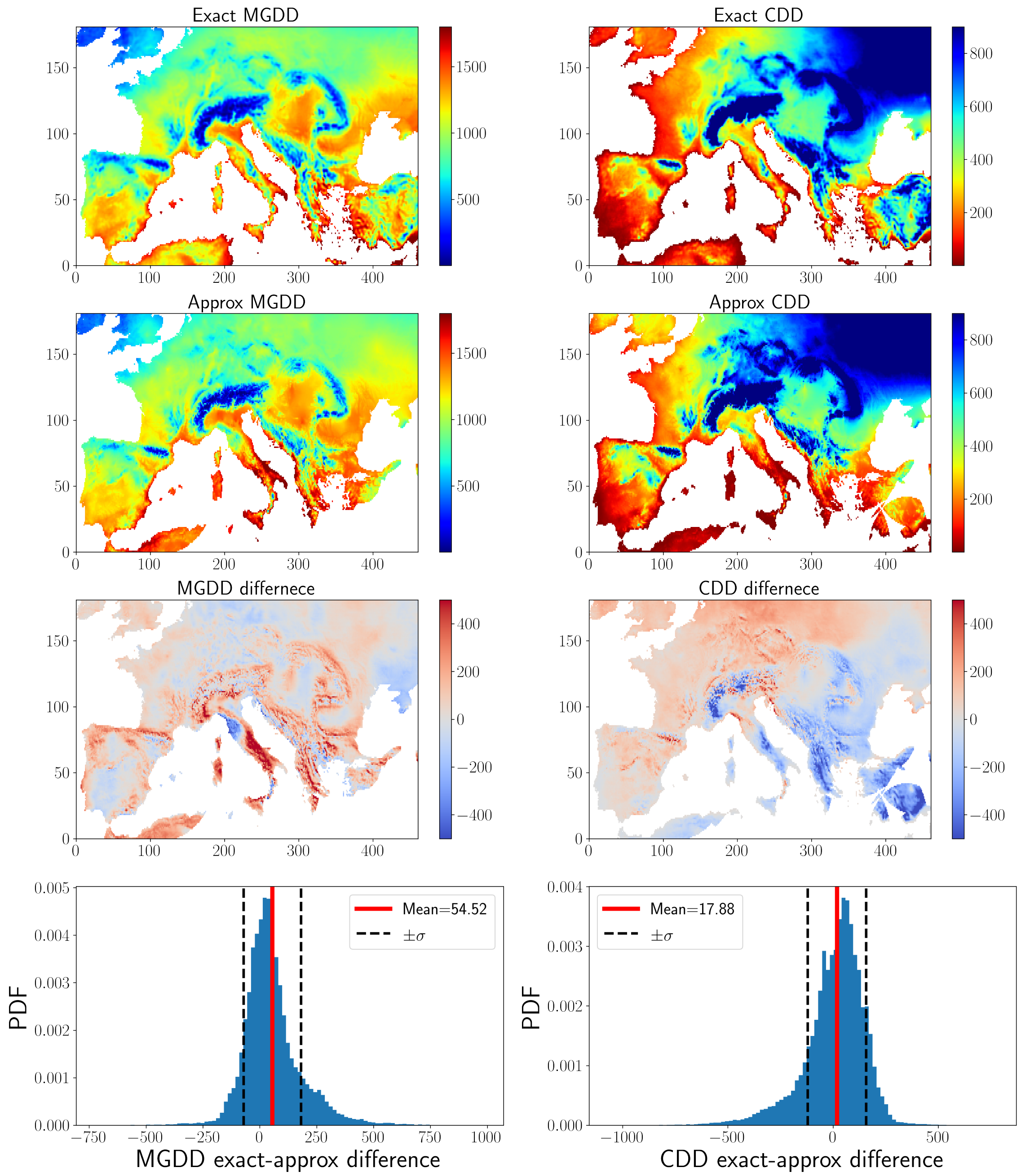


**Figure S7:** Differences in annual accumulated MGDD and CDD when using ERA5-Land dataset with hourly mean temperature and EOBS dataset with maximum and minimum daily temperatures.
